# A versatile and modular tetrode-based device for single-unit recordings in rodent *ex vivo* and *in vivo* acute preparations

**DOI:** 10.1101/2020.02.11.940809

**Authors:** Francisca Machado, Nuno Sousa, Patricia Monteiro, Luis Jacinto

## Abstract

The demand for affordable tools for recording extracellular activity and successfully isolating single units from different brain preparations has pushed researchers and companies to invest in developing and fabricating new recording devices. However, depending on the brain region of interest, experimental question or type of preparations, different devices are required thus adding substantial financial burden to laboratories. We have developed a simple and affordable tetrode-based device that allows interchangeable extracellular recordings of neural activity between *in vivo* and *ex vivo* preparations and can be easily implemented in all wet-bench laboratories. Spontaneous activity from several putative single neurons could be easily recorded and isolated by lowering the device into *ex vivo* cerebellum brain slices. The same device was also used *in vivo*, lowered into primary auditory cortex of adult anesthetized transgenic mice expressing channelrhodopsin in cortical neurons. Acoustic stimulation of the contralateral ear or direct laser optogenetic stimulation successfully evoked cortical activity at the recording site. Several isolated putative single neurons presented time-locked activity response to the different stimuli. In summary, we developed an affordable, versatile and modular device to facilitate tetrode extracellular recordings interchangeably between *in vivo* anaesthetized animals and *ex vivo* brain slice recordings.

**Highlights:** Developed a versatile and modular device to facilitate tetrode acute brain recordings interchangeably between *in vivo* and *ex vivo* preparations.

Conducted *ex vivo* extracellular recordings in acute cerebellar slices.

Conducted *in vivo* extracellular recordings in auditory cortex of anaesthetized mice.

Recorded and isolated multiple single units in both acute slices and anaesthetized mice recordings using the same device.

Device can be easily extended to accommodate optic fiber and cannula.

## 1. Introduction

Extracellular neuronal activity has been recorded and analyzed in neuroscience research for decades, providing insight into cell- and circuit-level brain computations [1], [2]. The ability to perform spike-sorting on extracellular activity recordings has been particularly useful since it allows the isolation of putative single neurons and the analysis of their activity/responses individually [3]. However, spike sorting is less commonly used in *ex vivo* brain slice recordings because typical single-wire pipette extracellular recordings are not ideal for this purpose and commercially available options are usually costly, namely multi-electrode arrays (MEA) [4], [5] and especially designed silicon probes [6]. Tetrodes are a simple and cost-effective approach that has been routinely used for chronic recordings of extracellular activity in freely-moving rodents [7]. Their long lasting success in neuroscience relies not only on their cost and easy implementation but also on their ability to facilitate spike sorting [7], [8]. Surprisingly, despite their success in freely-moving electrophysiology, tetrodes have been less frequently used in acute preparations including *in vivo* anesthetized animals and especially *ex vivo* slice preparations. One main factor limiting the use of tetrodes for the recording of extracellular spikes and subsequent unit isolation in acute preparations is the non-existence of commercially available devices that can simultaneously work as tetrode interface boards and structurally support/guide tetrodes into slices or whole-brains in anaesthetized animals.

Here, we propose a versatile and modular tetrode-based device that takes inspiration from probe design and freely-moving rodent tetrode microdrives, working for both *ex vivo* brain slices and *in vivo* anaesthetized recordings. The device parts can be easily produced by affordable and widespread technologies, many times in-lab, such as printed circuit board (PCB) manufacturing and 3D printing. Additionally, the device can be: quickly assembled; integrated with any amplifier/recording system already available at the lab for extracellular recordings and; used unlimited times, including inter-changeably between *ex vivo* and *in vivo* experiments, with multiple tetrodes and configurations. For *in vivo* preparations it can also be easily extended to be coupled with optic fiber or cannulas for optogenetic and pharmacological experiments. Using this device, we recorded spontaneous and opto- and sound-evoked neuronal activity in transgenic mice, both *in vivo* and *ex vivo*. In both configurations it was possible to reliably isolate several good quality single units from short recordings and analyze individual units’ responses to light and sound modulation. This device will facilitate a more widespread use of tetrodes for acute preparations in neuroscience experiments.

## 2. Materials and methods

### 2.1 Tetrode fabrication

Nichrome tetrodes (NiCr,12.5 µm diameter, RO-800 Hard PAC, Sandvik) were fabricated with standard methods, described elsewhere [9]. Briefly, a 50 cm long strand of insulated NiCr wire was folded and twisted by a motorized tetrode spinner (Tetrode Spinner 2.0, Neuralynx) while being tensioned. After the twisting procedure, the wires’ insulation was fused at 420°C with the aid of a heat gun and then carefully cut at the top and bottom. Tetrodes were then stored for later use.

### 2.2 Electrode interface board and tetrode guide structure

The proposed device consists of two parts: an electrode interface board (EIB) and a tetrode guide structure (Figure 1).

**Figure 1.**
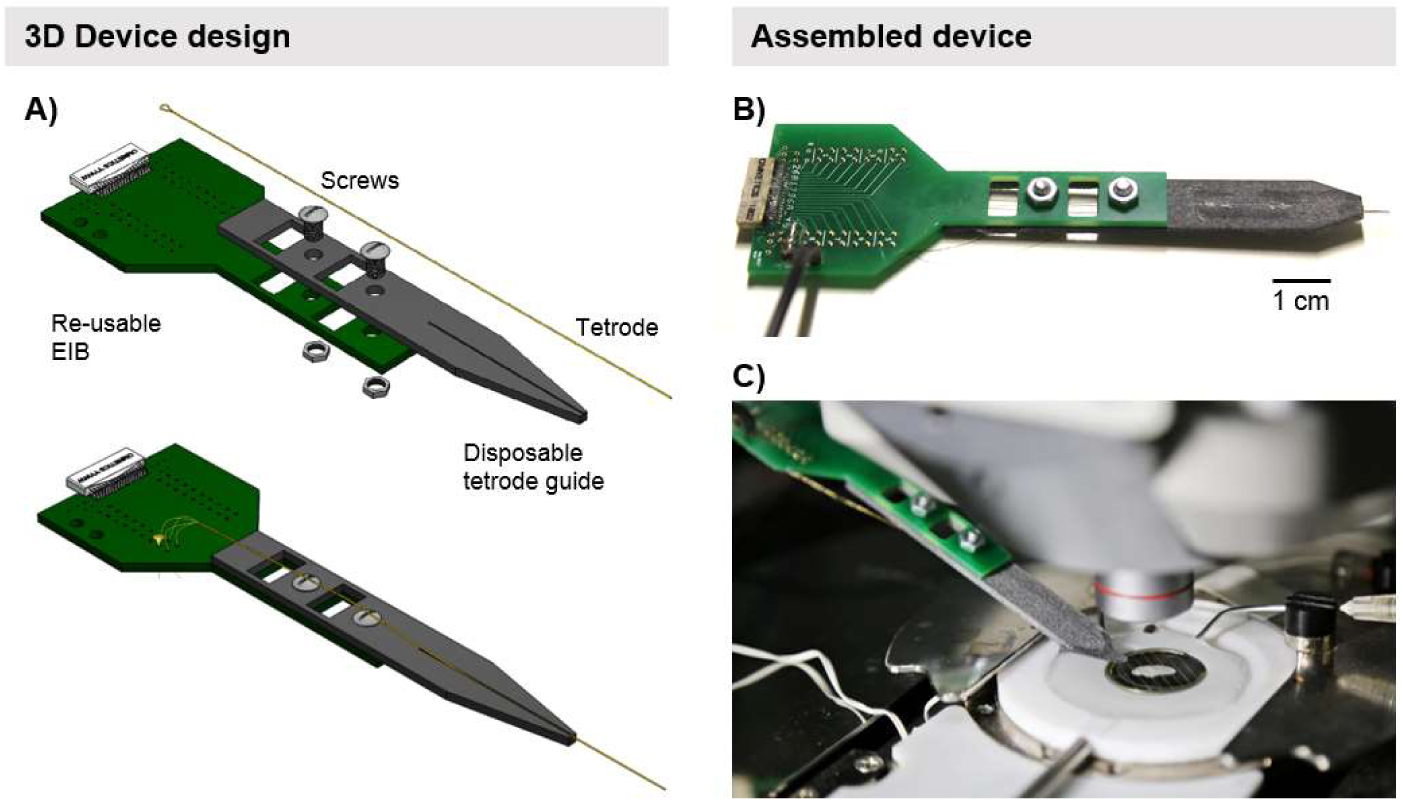
Tetrode device design overview. (A) 3D rendering of the device’s parts, including the electrode interface board (EIB) and the tetrode guide, separately (top) and fully assembled (bottom). (B) Photo of assembled device with loaded NiCr tetrodes and ground wire. (C) *Ex vivo* recording with device being held by micromanipulator and tetrodes guided into the slice inside the recording chamber.

The EIB is a printed circuit board (PCB) that allows the electrical connection of tetrode wires to gold coated vias (0.3 mm diameter) in the board by means of gold pins (small EIB pins, Neuralynx). These vias are routed to a female omnetics connector (A79024-001, Omnetics Connector Corporation) that mates with an appropriate headstage with male omnetics connector (which are used by many electrophysiology companies including Open Ephys, Neuralynx and Plexon, for example). There are 32 electrode vias (for 8 tetrodes) plus 2 ground and 2 reference vias. The 2 ground vias are shorted together. The EIB is a 2-layer PCB (10 mil trace/space, 1.6 mm thickness) and was manufactured with electroless nickel immersion gold (ENIG) surface finishing. The omnetics connector was soldered to the board by hot air soldering with solder paste (Leaded, No Clean Solder Paste, MG Chemicals). A 40 cm long 4-gauge isolated copper wire was looped through two drill holes on the PCB, designed for the purpose, and soldered to the ground vias.

The tetrode guide structure physically holds and guides the tetrodes into the tissue. This structure was 3D-printed in plastic (PA12 multijet fusion plastic; 70 × 10 × 1.5 mm) and has a single groove track that, depending on dimensions, can hold up to 8 tightly packed tetrodes lined in a single row. The support structure can be printed in any plastic material by any 3D printer with sufficient precision to print the tetrode guide track. The number, size and position of the tracks can also be easily adjusted by simple changes to the 3D part design file. The EIB and the tetrode guide structure were connected together by means of two 2mm diameter screws. There are two square holes on both parts that line up and allow passing of tetrode wires from one side of the board to the other, if necessary.

Tetrodes were placed in the guide track with the aid of ceramic coated tweezers (11252-50, Fine Science Tools) under a microscope (SZ51, Olympus) and glued in place with 5-minute epoxy (Devcon). Each tetrode wire was then routed to an electrode via in the EIB and secured with a gold pin. Tightly pressing the gold pin against the electrode wire inside the via, removes the electrode’s insulation and allows electrical connection. Figure 1B shows a fully assembled device with 8 NiCr tetrodes loaded and connected to the EIB.

### 2.3 Tetrode electrodeposition

After gluing the tetrodes to the guide structure and physically connecting each tetrode wire to the electrode vias in the EIB, tetrodes’ tips were cut with sharp stainless-steel scissors (14568-09, Fine Science Tools) and electroplated. The plating solution consisted of 5% gold non-cyanide solution (Sifco 5355 Gold Plating Solution, Neuralynx) with 1% Poly(ethylene glycol) (PEG) addictive (BioUltra, 8,000, Sigma-Aldrich) mixed in a 75% v/v PEG to gold ratio. This solution has been shown to promote a large surface area with low impedance [10]. Electrode impedance was lowered to 150-200 KOhm in successive steps of electrodeposition at −0.05 µA with the aid of NanoZ (White Matter LLC).

### 2.3 Ex-vivo and in-vivo electrophysiological recordings

#### 2.3.1 Animals

*Emx1-Cre:Ai27D* or *Pvalb-Cre:Ai27D* male mice were originally purchase from JAX (stocks #008069, #005628, #012567) [11]–[13]. All experiments were conducted in accordance with European Union Directive 2016/63/EU and the Portuguese regulations and laws on the protection of animals used for scientific purposes (DL N° 113/2013). This study was approved by the Ethics Subcommittee for the Life Sciences and Health of University of Minho (ID: SECVS 01/18) and the Portuguese Veterinary General Direction (ID: DGAV 8519).

#### 2.3.2 Ex vivo cerebellar electrophysiological recordings

##### 2.3.2.1 Ex vivo slice preparation and recordings

Briefly, animals were deeply anaesthetized with avertin (0.5 mg/g body weight; tribromoethanol; 20 mg/mL; Sigma–Aldrich) by intraperitoneal injection and decapitated. The brain was quickly removed to prepare 300 μm cerebellum parasagittal slices according to the protocol of [14]. All recordings were performed at 34 °C in artificial cerebrospinal fluid gassed with 5% CO_2_/95% O_2_. The tetrode device containing 8 NiCr electroplated tetrodes was secured to a 3-axis micromanipulator (PatchStar, Scientifica) and connected to a 32-channel headstage (RHD2132, Intan). The tetrodes were then guided into the slice with the micromanipulator under visual inspection of a microscope (Figure 1C) and placed close to the Purkinje cell layer at cerebellum lobule IV. Extracellular activity was recorded with an Open Ephys acquisition system (Open Ephys) [15] connected to the headstage by serial peripheral interface (SPI) cables (Intan). Signals were acquired at 30Ks/s.

#### 2.3.3 In vivo auditory cortex electrophysiological recordings

##### 2.3.3.1 Surgical procedure and in vivo extracellular recordings

Mice were anesthetized by intraperitoneal injection mix of ketamine (75mg/Kg) and medetomidine (1mg/Kg) and positioned in a stereotaxic frame (Stoelting). Dorsal skull was exposed and the animal was rotated 90° degrees to facilitate removal of the temporal muscle and access to the primary auditory cortex (A1). A 1.5 × 1.5 mm area above A1 centered at −2.5 mm AP and 4.0 mm ML from bregma [16] was opened and the dura removed under a microscope (S6, Leica Mycrosystems). The tetrode device containing 8 NiCr electroplated tetrodes was attached to a micrometric stereotaxic arm (1760, Kopf Instruments) and connected to an headstage (RHD2132, Intan). Tetrodes were lowered into the brain, through the skull opening, to −0.6 mm (DV) from the brain surface. A stainless-steel screw electrode (E363-20-2.4-03, Plastics One) secured in a burr hole at the back of the skull served as ground. The surgical procedure and the following stimulation experiments were conducted on a custom-made anti-vibration table inside a double-walled sound-proof chamber. Extracellular signals were acquired with an Open Ephys acquisition system at 30 Ks/s.

##### 2.3.3.2 Evoked auditory response

To evoke responses from A1 neurons, tones of variable frequency (8, 12, 20, 24, 28 and 32 KHz, 25 ms duration, 975 ms intertrial interval) and variable sound pressure level (50, 60, 70 and 80 dB SPL) were pseudo-randomly delivered from a free field electrostatic speaker (ES1, TDT) positioned 10 cm away from the mouse’s contralateral ear. The speaker was driven by a speaker driver (ED1, Tucker-Davis Technologies) and tones were generated by an high-sampling rate sound card (HARP, Champalimaud Foundation) controlled by Bonsai [17].

##### 2.3.3.3 Optogenetic stimulation

To evoke neuronal activity of A1 neurons, the surface of the exposed brain was illuminated with blue light pulses (473 nm wavelength, 10 Hz with 30 ms pulse width, 5s On, 55s Off). Light was delivered by an optical fiber (200 μm diameter) connected to a fiber-coupled DPSS laser source (CNI). Light pulses were generated by a waveform function generator (DG1022, Rigol) connected to the laser source. The optical fiber was held by 3-axis micromanipulator (LBM-2025-00, Scientifica) and placed 5 mm above brain surface.

#### 2.3.4 Signal processing and spike sorting

Signals were analyzed with custom-written Matlab (Mathworks) code. Both *ex vivo* and *in vivo* recordings were filtered between 0.6 and 6 KHz. Spikes were detected using a variable amplitude threshold that was a multiple (4 to 6 times) of the median from an estimate of the background noise’s standard deviation as in [18]. Spike sorting was achieved by performing weighted principal component analysis (wPCA) on the waveforms of detected spikes followed by Gaussian Mixture Model (GMM) unsupervised clustering of the first principal components of pairs of electrodes from the same tetrode (Figure 2B) [19]. In a final step, clusters were manually split or merged based on visual inspection of cluster waveforms and their inter-spike interval histograms and auto-correlograms.

**Figure 2.**
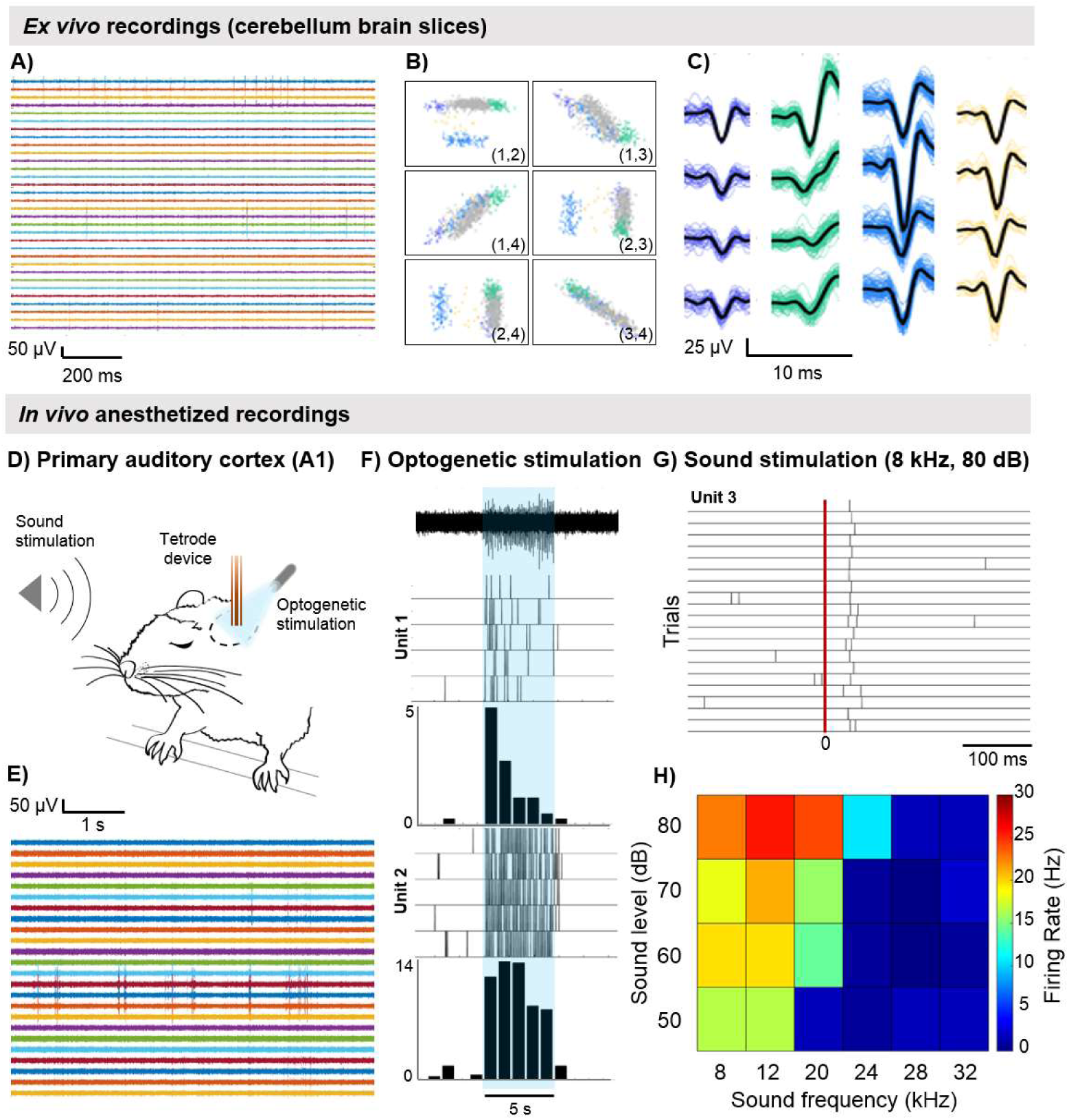
*Ex vivo* and *in vivo* electrophysiological recordings. (A) Extracellular signals recorded on 8 tetrodes from a mouse cerebellum brain slice (signals band-passed filtered between 0.6 and 6 KHz). (B) Cluster isolation for one tetrode using PCA and GMM based spike sorting. Each scatterplot shows the projections of two pairs of tetrode’s channel waveforms onto the first principal component. Colored points correspond to isolated single units (matching colors with respective waveforms on right). Grey points correspond to multi-unit activity which was not possible to isolate into single units. (C) Sample waveforms of four isolated single units from one tetrode (average waveform in black). (D) Representative drawing of the *in vivo* experiment combining tetrode recordings with optogenetic and sound stimulation in primary auditory cortex (A1) of an anesthetized mouse. (E) Extracellular signals recorded on 6 tetrodes from A1 (signals band-passed filtered between 0.6 and 6 KHz). (F) Evoked activity of isolated single units in A1 by optogenetic stimulation. Extracellular activity recorded during one trial of light stimulation (light stimulation period in blue, corresponding to 5 seconds) (top); Raster plots and average peri-stimulus time histograms of two isolated single units over five trials. (G-H) Evoked activity of one isolated single unit in A1 by sound stimulation. (G) Raster plot of one isolated single unit responding to 8 KHz tone stimuli over 20 presentations (red line marks tone presentation). (H) Tuning heatmap of the same unit showing average firing rate across the different sound frequencies and sound levels presented during the evoked auditory response protocol.

## 3. Results

The device was assembled and ready for recordings in under 2 hours including 1 hour for fabrication of eight tetrodes and 1 hour for assembly, tetrode loading and electroplating. The same tetrode tips were cut fresh and electroplated between the *ex vivo* and *in vivo* experiments.

In the *ex vivo* brain slice experiment (Figure 2A-C) we were able to isolate nine putative single neurons from eight tetrodes positioned at the Purkinje cell layer (cerebellum lobule IV) during a 30 minute recording of spontaneous activity. Figure 2C shows 4 units isolated from a single tetrode wire.

In the *in vivo* experiment (Figure 2D-H) several putative neurons were isolated from six tetrodes during a 30 minute recording from the primary auditory cortex in an anesthetized mouse. Additionally, by identifying putative neurons through spike sorting, we were able to analyze single-units’ responses to specific light and sound stimuli (Figure 2F-H).

## 4. Discussion

This manuscript describes the design and implementation of an affordable, versatile and modular tetrode-based device that allows extracellular recordings in both *ex vivo* and *in vivo* preparations. The same device can be used in both preparations without any changes in configuration or parts, as performed here. Its probe-like design allows easy positioning in micromanipulators for *ex vivo* slice recordings and in stereotaxic frames for *in vivo* recordings. The proposed device is solely based on parts that can be easily produced in-lab or sourced for a low cost. This is especially important considering the cost of MEAs and silicon probes, which might be prohibitive for many labs especially those running experiments demanding constant re-use.

The EIB was inspired by EIBs from microdrive systems for tetrode positioning in freely-moving animals [20]–[22] where the easiness of assembly must be combined with quick and non-damaging part recovery. PCB manufacturing is currently a widespread service accessible at a low cost. The present EIB was designed to match headstages with male omnetics connectors which are used by many traditional electrophysiology systems such as Open Ephys (used in the experiments described here), Neuralynx and Plexon; but can be easily changed and produced to accommodate other types of connectors for other headstages.

Similarly, 3D printing has also become an affordable technology and most labs now have access to it in-house. The tetrode guide structure design was inspired by silicon probe tip’s design and can be printed in virtually any commercial or in-house 3D printer, contingent to printer’s precision and the desired guide track’s dimensions. Since 3D printing allows easy customization, the proposed design for the guide structure can be easily changed by simple alterations to the 3D part design file to accommodate several configurations, from single tetrode tracks to multiple tracks in the same or different locations. The possibility of changing the track dimensions and position allows versatile tetrode positioning – for example, for targeting different brain structures simultaneously – and the use of tetrode wires of different diameters for different experimental demands. The guide design presented here can also be modified to accommodate optic fiber tracks for optogenetic experiments or cannula for pharmacological experiments. Despite this versatility, and although tetrodes can be cut with precision scissors and lined up in a row with micrometric precision under a microscope, when compared with MEAs and silicon probes, their real geometry is not fixed/known which requires, for example, additional modeling for current source density analysis studies [23].

The whole device was designed for constant re-use with minimal cost and minimal damage to the parts upon recovery. In fact, the same tetrodes glued to the guide structure and connected to the EIB can be used for several sequential recordings across different days as long as tetrode tips are cut fresh and electroplated between experiments (and as long as the tetrodes are of sufficient length for the intended target). Nevertheless, the whole device can be quickly recovered for a new experiment. The EIB can be re-used by removing the gold pins from the vias and discarding the tetrodes. The same gold pins can be re-used for the next application. The tetrode guide can be recovered by immersion in 70% alcohol solution overnight to remove the epoxy, or it can also be discarded between experiments given that it can be printed effortlessly.

The affordability and easiness of manufacturing and assembly of the proposed device will allow a more widespread use of tetrodes for acute extracellular recordings and unit isolation in *ex vivo* and *in vivo* preparations.

## Acknowledgements

The authors would like to acknowledge Diana Rodrigues for helping preparing *ex vivo* brain slices and Margarida Gonçalves for helping with *in vivo* A1 surgeries. This work was supported by Calouste Gulbenkian Foundation (grant number P-139977); Society in Science, The Branco Weiss fellowship, administered by Eidgenössische Technische Hochschule (ETH) Zürich; the European Molecular Biology Organization (EMBO) Long-Term Fellowship (ALTF 89-2016 to P.M.) and FCT (grant number PTDC/MED-NEU/28073/2017, POCI-01-0145-FEDER-028073). This work was also funded by FEDER through the Competitiveness Factors Operational Programme (COMPETE), by National funds through the Foundation for Science and Technology (FCT) under the scope of the project UID/Multi/50026; and by the project NORTE-01-0145-FEDER-000013, supported by the Northern Portugal Regional Operational Programme (NORTE 2020), under the Portugal 2020 Partnership Agreement, through the European Regional Development Fund (FEDER).

